# A rapid and efficient *in vivo* inoculation method for introducing tree stem canker pathogens onto leaves: Suitable for large-scale assessment of resistance in poplar breeding progeny

**DOI:** 10.1101/2024.03.13.584916

**Authors:** Zheng Li, Bingyu Zhang, Yuchen Fu, Yutian Suo, Yinan Zhang, Jinxia Feng, Long Pan, Wanna Shen, Huixiang Liu, Xiaohua Su, Jiaping Zhao

**Affiliations:** State Key Laboratory of Tree Genetics and Breeding, Institute of Ecological Conservation and Restoration, Chinese Academy of Forestry, Beijing, China; State Key Laboratory of Tree Genetics and Breeding, Research Institute of Forestry, Chinese Academy of Forestry, Beijing, China; Shandong Research Center for Forestry Harmful Biological Control Engineering and Technology, College of Plant Protection, Shandong Agricultural University, Taian, China

**Keywords:** poplar hybrid breeding, tree diseases, stem canker, *in vivo* leaf inoculation, *Valsa sordida*

## Abstract

Hybrid breeding, a direct and efficient strategy for disease control and management in tree species, is currently limited by the selection method of resist clones: the “*in vitro* stem segment inoculation method.” This method, constrained by the availability of inoculating materials, cannot rapidly, efficiently, and cost-effectively screen the resistance of all hybrid clones. To overcome these limitations, we introduce a novel pathogen inoculation method for the resistance assessment of hybrid clones in the poplar-*Valsa sordida* pathosystem. This method involves inoculating the stem canker pathogen on the host leaf, a unique and promising approach we have successfully validated.

Results showed that stem canker pathogen *V. sordida* induced the extended necrotic lesion and even induced the formation of pycnidium structure and conidia on the leaf surface five days after mycelium inoculation; 1) the upper 5-7^th^ leaves exhibited higher resistance than the middle 18-20^th^ leaves; 2) the shading conditions induced more severe symptoms on the leaves than lighting conditions; 3) the juvenile mycelium inoculums (4-day-cultured) were more susceptible to poplar leaves than the old ones (7-day-cultured). Our results demonstrate the robustness of the “*in vivo* leaf inoculation method” in revealing the resistance differentiation in poplar hybrid clones. According to the leaf necrotic area disease index, we divided these poplar clones into seven different resistance groups. The resistance assessed by leaf assessment was validated in 15 selected poplar clones using the “*in vitro* stem segment inoculation method.” Results showed that the effectiveness of these two methods was consistent. Moreover, the leaf inoculation method can be used to detect the pathogenicity diversity of the pathogen population of tree species. Compared to the conventional “*in vitro* stem segment inoculation method,” the leaf method has the advantages of abundant inoculation materials, easy operation, rapid disease onset, and almost no adverse effect on the host. It is particularly suitable for the resistance screening of all progeny and the early (seedling) phenotypic selection of resistant poplar clones in poplar stem disease resistance breeding. The “*in vivo* leaf inoculation method” holds significant promise in poplar breeding, tree pathology, and molecular biology research on tree stem diseases.

## 1 Introduction

Poplar species are frequently subjected to attacks by various pests, including two prominent stem canker pathogens: *Botryosphaeria dothidea* (Moug. ex Fr.) Ces. & De Not. (anamorph *Fusicoccum aesculi* Corda) and *Valsa sordida* Nitschke (anamorph *Cytospora chrysosperma* (Pers.) Fr.). These pathogens are major contributors to forest diseases in China, capable of devastating poplar seedlings across plantations or causing significant damage to mature poplar forests (Zhang & Luo, 2003). The most direct and effective method for managing and controlling forestry diseases involves hybridization breeding. Developing resistant cultivars through hybridization, genetic modification, or gene editing is recognized as the most efficient, cost-effective, and environmentally sustainable strategy to combat plant diseases, such as powdery mildew (Yin & Qiu, 2019).

To effectively breed resistant clones against plant diseases, it is imperative to precisely and rapidly determine disease phenotypes (pathophenotypes)—including disease incidence rate (IR), onset time, necrotic area (NA, defined as the average area size of necrotic spots), and disease index (DI). This constitutes a critical challenge in pathology that impacts breeding efficiency. Three primary methodologies are employed to ascertain the disease phenotypes for breeding poplar canker: field investigation assays, callus inoculation (Zhang *et al*., 1989), and *in vitro* stem segment inoculation. The *in vitro* stem segment inoculation method, frequently utilized (Shi *et al*., 2014; Zhang *et al*., 2017), involves using eight to ten stem segments, each 30-40 cm in length, as inoculation materials. Each segment is inoculated at five points, with IR monitored and recorded every five days over a 60-day period to determine the DI. The poplar segments typically originate from either 1- or 2-year-old saplings or mature poplar trees that have been cultivated in the field for over four years, belonging to various poplar hybridization clones. However, this method also has its drawbacks: it is time-consuming, requires significant experimental space, and thus incurs considerable costs in terms of breeding time, land, labor, and expenditure.

In contrast to stems, leaf tissue is abundant and rapidly growing, making it an excellent candidate for assessing plant resistance to stem canker disease. Prior to conducting an *in vivo* leaf inoculation to evaluate host resistance, it is crucial to verify the presence of stem canker pathogens in poplar leaves. The initial and most critical step in disease onset involves the invasion and colonization of pathogens in host tissues. Thus, confirming the presence of stem canker pathogens in poplar leaves is essential before implementing this method. *B. dothidea* can cause canker disease on stems and branches of plants but may also reside latently as endophytic pathogens within leaves, fruits, and other plant parts (Marsberg *et al*., 2017). Previous research employed excised leaves of *Malus spectabilis* to evaluate the pathogenicity of mutant isolates of *B. dothidea* (Zheng *et al*., 2017). Similarly, the mycelium and conidial suspension of *V. ceratosperma*—a stem canker pathogen of apple trees—can induce disease symptoms on excised leaves, shoots, and fruits of apple trees (Wei *et al*., 2010; Xie, 2019). Additionally, *Septoria musiva* Peck, a significant fungal pathogen in hybrid poplars across the north-central and northeastern regions of North America, causes both leaf spot and stem canker diseases in hybrid poplars (Feau *et al*., 2010; Dunnell & LeBoldus, 2017). Based on this research, inoculating stem canker pathogens may induce fungal diseases in leaf tissues. Zhang *et al*. (2018) suggested using excised leaves to assess the pathogenicity of rice blight pathogens. Similarly, inoculating isolated leaves and shoots of *V. ceratosperma* provides a rapid and precise method to assess the pathogenicity of the stem canker pathogen in apple trees (Wei *et al*., 2010). However, no specific protocol for this method is proposed in this work, nor is its application in resistance screening for hybrid clones in tree stem disease breeding discussed in other research. Consequently, the acquisition of pathophenotypes and the selection of resistant clones remain technical challenges in the hybrid breeding of poplar stem canker diseases.

This study introduces a novel method for assessing resistance to stem canker pathogens in poplar clones through leaf inoculation. It evaluates the impact of leaf developmental stage (specifically, the position of the leaf on the branch), light conditions, and the cultivation period of the fungal pathogen on the resistance of poplars, utilizing the poplar/*V. sordida* disease system. Resistance to *V. sordida* is measured within a small, randomly selected cohort of hybrid poplar clones, and these findings are confirmed through stem segment inoculation techniques. Additionally, the research explores the diversity of pathogenicity among various *B. dothidea* isolates in poplar. The ultimate goal is to develop a rapid, effective, and cost-efficient method for inoculating stem canker pathogens, along with a new approach to hybrid breeding for managing canker diseases in poplars and other tree species.

## 2 Materials and methods

### 2.1 Fungal and plant materials

This study employed the already identified poplar canker pathogen *V. sordida* isolate CZC (NCBI accession number: MK994101 for rRNA-ITS and MN025273 for EF1α gene) along with 13 isolates of the poplar blister canker pathogen *B. dothidea* (Table 1) (Xing *et al*., 2020; Wang, 2013).

**Table 1.** Hosts and locations of fungal isolates used in this study.

All fungal isolates were maintained at the Laboratory of Plant Physiology within Institute of Ecological Conservation and Restoration, Chinese Academy of Forestry in Beijing. The isolates were cultured on potato dextrose agar (PDA) medium (20% potato, 2% dextrose, 2% agar; pH 6.0) at 28°C under dark conditions for 4 or 7 days after inoculation (DAI). Subsequently, the mycelium-covered medium was cut into square pieces, each approximately 1.0-1.2 cm in length.

For the plant materials, the study utilized 1- and 2-year-old saplings of *Populus euramericana* cv. ’Bofeng 3’ (Huang *et al*., 2014), 6-year-old *P. alba* var. *pyramidalis* saplings, and 2-year-old saplings of the hybrid clones (*P. deltoides* × *P. deltoides*, 48 genotypes). These saplings were grown either in the greenhouse or in the experimental fields of CAF, ensured to be free from pest infestation on leaves and stems and adequately irrigated throughout the duration of the experiments. Prior to inoculation, three to four mature and adjacent leaves were washed using sterilized water and subsequently sterilized with 75% alcohol (vol/vol) (Figure 1A). After air-drying, all sterilized leaves were marked for subsequent inoculation.

**Figure 1:**
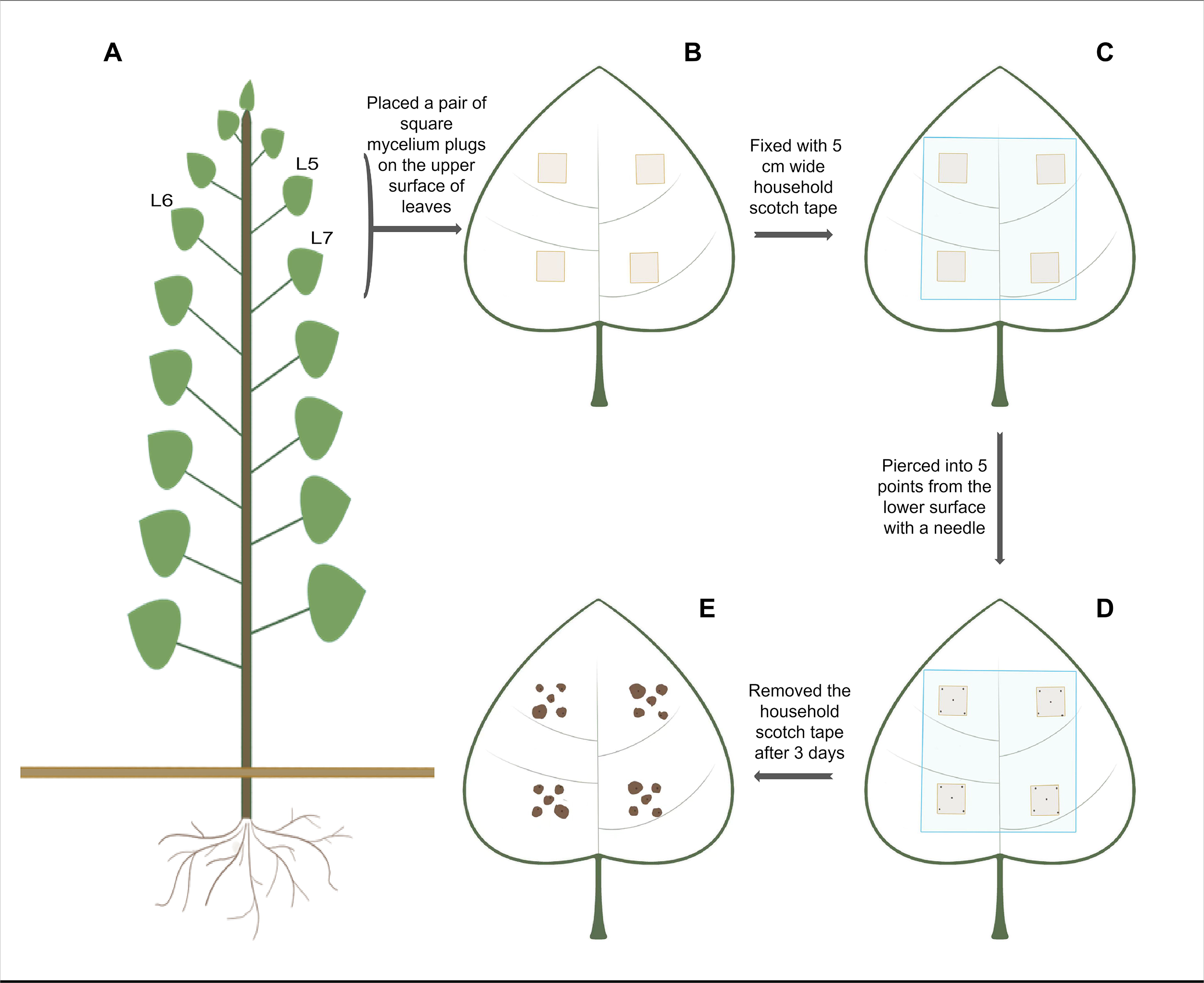
Schematic diagrams of *in vivo* leaf inoculation methods used in this study.

### 2.2 Inoculation methods

Depending on the leaf size, either two square pieces of PDA medium covered with mycelium (for smaller leaves with a width of less than 10 cm) or four square pieces (for larger leaves with a width exceeding 10 cm) were inoculated onto the selected leaves. The inoculation sites were strategically placed on the upper surface of the poplar leaves, approximately 1-3 cm from the central veins, ensuring that the secondary veins were not obscured by the mycelium pieces (Figure 1B). The mycelium was oriented to face the leaf surface. Additionally, for control purposes, one leaf from each poplar genotype adjacent to the pathogen-inoculated leaves was inoculated with 2% water agar (WA) square pieces. Following pathogen inoculation, the leaves were covered and sealed with 5 cm wide household scotch tape to secure the mycelium or WA pieces and prevent moisture loss (Figure 1C). The leaves were then pierced at five points on the lower surface with a needle, either at the center of the inoculated mycelium or WA pieces or near the corners of the squares, approximately 1-2 mm from the corners (Figure 1D). Post-inoculation, the integrity of the scotch tape seals and the development of disease symptoms on the inoculated leaves were monitored and documented daily (Figure 1E). At 5 DAI, all inoculated leaves were harvested from the poplar branches or twigs, placed in dark plastic bags, and stored at 4°C. Upon completion of the experiment, the leaves were brought to the laboratory; after removing the scotch tapes, mycelium, and WA pieces, the leaves were photographed or scanned as JPEG images (minimum resolution of 1500 pixels). The necrotic spots around the pricking sites and the area of these spots were identified automatically using ImageJ software (https://imagej.nih.gov/ij/) or manually adjusted when necessary. Finally, the results were compiled in Microsoft Excel.

In this study, the criteria for the occurrence of leaf disease were established based on the lesion shape, color, size, and the presence of hypha-like or pycnidia-like structures around the pricking sites inoculated with a pathogen (or WA). In treatments where leaves were inoculated, disease symptoms manifested as lesion spots around the pricking sites, expanding to exceed 2.0 mm^2^ within five days; additionally, grey or dark hypha-like structures, or occasionally pycnidia-like structures, were observed on the surface of the necrotic spots. In contrast, mechanically induced leaf lesions in the mock inoculation treatment remained brown, rarely expanded within five days, and maintained an average size of approximately 1.5 mm^2^, with no hypha-like structures forming. For simplicity, the value of the lesion areas on the inoculated leaves was set at A=2.0 mm^2^ as the quantitative criterion to determine the occurrence of leaf disease (Table 2). If A > 2.0 mm^2^, the pricking sites were considered diseased, and vice versa. Disease incidence rate (IR) represents the proportion of inoculation sites on poplar leaves that develop necrotic areas greater than 2.0 mm^2^ five days post- inoculation. Disease index (DI) is a quantitative measure that assesses and describes the incidence and severity of disease on the leaves, aiding in a more accurate understanding of the disease conditions on the foliage. Based on the lesion area values at each pricking site, the DI for each poplar clone was calculated, and a ranking level of disease development was proposed (Table 2).

**Table 2.** Severity grading of necrotic symptoms induced by canker pathogens on leaves

The incidence of disease and DI were calculated using the following formulas:

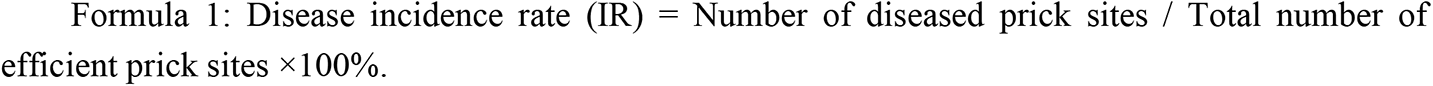

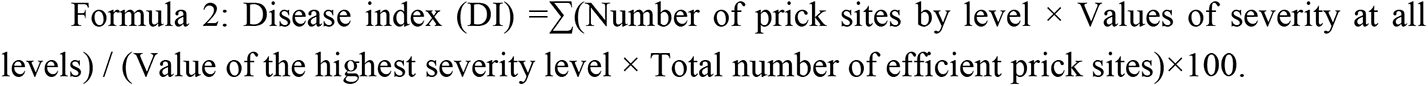

### 2.3 Comparison of resistance to canker pathogen across different stages of leaves

Square mycelium plugs of *V. sordida* isolate CZC, cultured in darkness for seven days, were inoculated onto the 5^th^ to 7^th^ newly matured leaves (from the top to the bottom of the poplar saplings, the same applies from now on) and the 18^th^ to 20^th^ mature leaves of three *P. euramericana* cv. ’Bofeng 3’ saplings. Each leaf received two square mycelium plugs. In total, 120 pathogen-inoculated pricking sites were established, with 60 on the upper leaves and 60 on the lower leaves. Additionally, WA plugs were used to inoculate another sapling as a control, creating 40 pricking sites. The details of the inoculation method are described in the Inoculation Methods section.

### 2.4 Comparison of resistance between light-exposed and shaded leaves

The experiment evaluated the resistance of leaves from 6-year-old *P. alba var. pyramidalis* branches inoculated with the *V. sordida* isolate CZC, which was cultivated in darkness at 28°C for seven days. The study involved three replicates of poplar, each with two branches—one exposed to light and one shaded. The base diameters of the branches were identical (1.42±0.14 cm for light- exposed branches and 1.45±0.17 cm for shaded branches; n=3 for each). The 5^th^ to 7^th^ newly matured leaves on each branch were inoculated with two square plugs of pathogen mycelium, resulting in a total of 120 pathogen-inoculated pricking sites (60 for the shaded group and 60 for the light-exposed group). Additionally, two branches from both the light-exposed and shaded groups were inoculated with WA plugs as controls.

### 2.5 Comparison of the pathogenicity of canker pathogen across different culture durations

To assess the effect of culture duration on the pathogenicity of *V. sordida* isolate CZC, this pathogen was cultivated in darkness for both 4 and 7 days before being inoculated onto the leaves of *P. euramericana* cv. ’Bofeng 3’. In this study, six poplar saplings were divided into two groups: one inoculated with the pathogen cultured for 4 days (D4) and another with the pathogen cultured for 7 days (D7). Two square mycelium plugs were used to inoculate the 5^th^ to 7^th^ newly matured leaves. A total of 60 pathogen-inoculated pricking sites were created for each group, and 30 pricking sites served as the control.

### 2.6 *In vivo* assessment of resistance to *Valsa* canker in hybrid poplar clones using leaf inoculation method

The *Valsa* pathogen isolate CZC was cultured on PDA medium in darkness at 28°C for seven days. The 5^th^ to 7^th^ newly matured leaves of one or two hybrid poplar clones were inoculated with two to four square mycelium plugs. Three leaves per clone were inoculated with pathogen mycelium plugs, and one leaf per clone was inoculated with WA plugs as controls. After inoculating all 48 hybrid clones, the leaves were immediately pricked with needles. In total, pathogen plugs were inoculated at either 30 or 60 pricking sites, and WA plugs at 30 pricking sites for controls. Disease symptoms were observed and photographed at 5 DAI. The symptoms and NAs around the pricking sites were used to determine the IR and DI.

### 2.7 *In vitro* validation of poplar resistance to *Valsa* canker pathogen using stem segments

To validate the effectiveness of the leaf inoculation method in assessing resistance to the stem canker pathogen, the responses of 15 *P. deltoides* × *P. deltoides* hybrid clones (5 resistant/highly resistant, 5 non-resistant/non-susceptible, and 5 susceptible/highly susceptible to the canker pathogen, as determined by *in vivo* leaf inoculation described in Section 2.6) were evaluated using an excised stem segment inoculation method. This study utilized ten dormant poplar stem segments (2-3 cm in diameter, 25-30 cm in length) sourced from the base region (50-60 cm) of 1-year-old branches from the selected poplar clones (Figure 2A). Before inoculation, these segments were thoroughly washed with sterilized water, sterilized with 75% alcohol, and the upper end was treated with plant callus ointment—a commercial product that promotes callus regeneration around wounds—once dry (Figure 2B). The segments were soaked in water for three days at room temperature before being inoculated with *V. sordida* isolate CZC (cultured on 2% PDA at 28°C for seven days) (Figure 2C). Zhang (2017) detailed an inoculation technique in which each stem segment was inoculated with five mycelium plugs, each 0.5 cm in diameter, spaced 3-4 cm apart, starting from 10 cm above the base of the segment and extending to the upper end. All inoculation sites were covered with household PE film to retain moisture (Figure 2D). Each site was pricked in a regular triangular pattern using a needle, with side lengths of approximately 1-2 mm (Figure 2F). After inoculation, the stem segments were maintained in water. In this research, ten segments underwent inoculation, with two serving as mock-inoculated controls.

**Figure 2:**
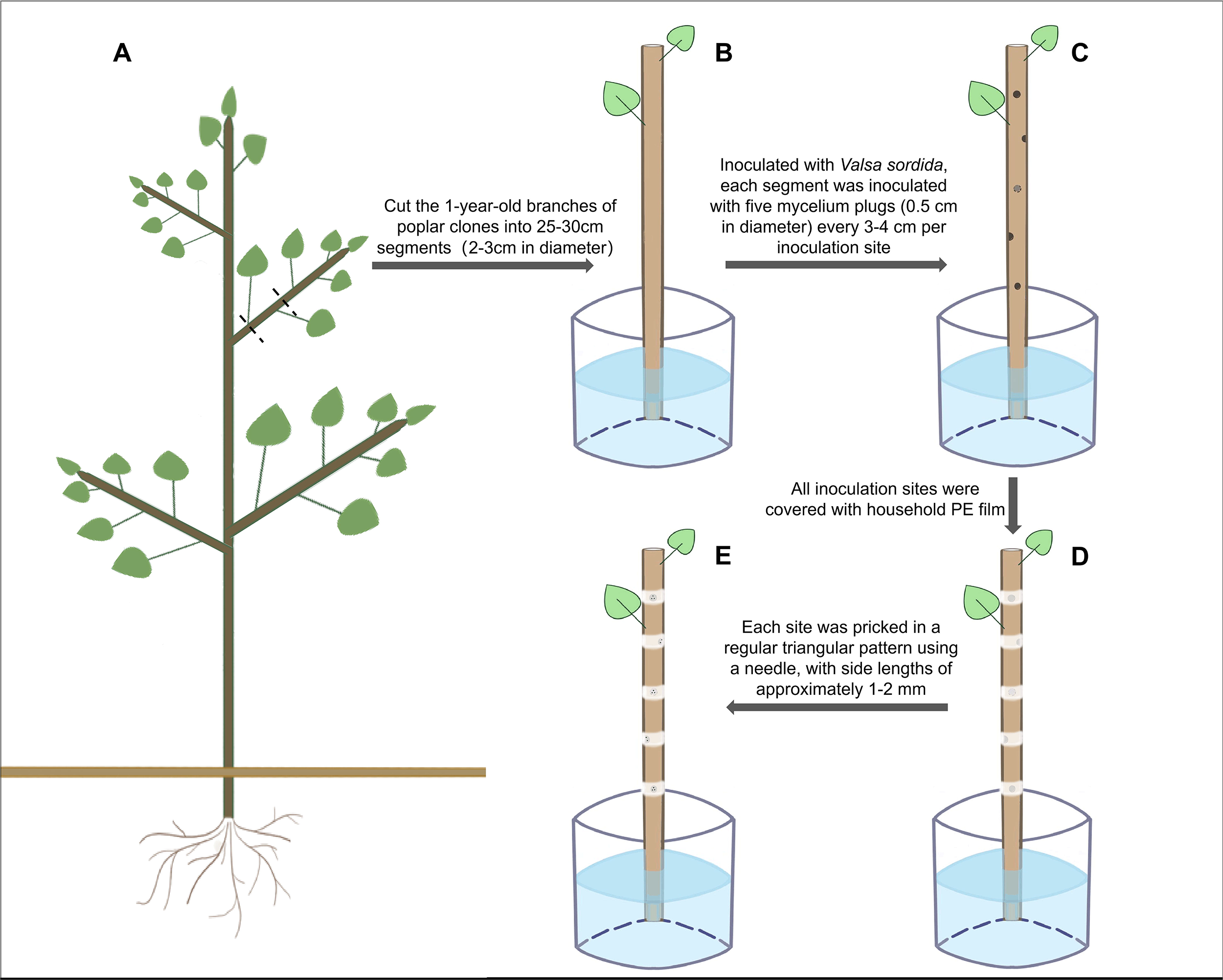
Schematic diagrams of *in vitro* stem segments inoculation methods used in this study.

Disease progression at the inoculation sites was monitored and recorded based on the appearance of necrotic tissue every five days. The necrotic sites were marked, and the total number of disease sites for each poplar clone was recorded until 40 days post-inoculation (Yang *et al*., 1989; Shi, 2014). Disease severity for each poplar clone was ranked according to the onset time of symptoms (Table S1) (Yang *et al*., 1989; Zhang *et al*., 1989), and the DI was calculated using Formula 2. At the final observation, the longitudinal extension length (a, in mm) and the horizontal length (b, in mm) of each lesion were measured. Given that the lesion shapes were predominantly subelliptical, the NA of the disease spots was calculated using the formula for the area of an ellipse. The severity of the disease was ranked according to the criteria in Table S2, and the DI was calculated using Formula 2.

### 2.8 Determination of pathogenicity of stem canker pathogens using *in vivo* leaf inoculation method

This study assessed the pathogenicity of 12 *B. dothidea* isolates sourced from various regions and hosts via the leaf inoculation method. The 5^th^ to 7^th^ newly matured leaves on 12 different twigs from a 2-year-old Populus ’Bofeng 3’ hybrid clone were inoculated with these fungal isolates. Each isolate was applied to three leaves using either two or four square mycelium plugs. Subsequent to needle piercing, either 30 or 60 pathogen-inoculated pricking sites were established for each isolate. Additionally, the leaves of two other twigs were inoculated with WA plugs and similarly needle pierced to serve as control treatments, denoted as Ctrl-1 and Ctrl-2. The leaf inoculation technique employed was detailed in the Inoculation Methods section. Five days post-inoculation, the symptoms were observed, captured in photographs, and analyzed using ImageJ software, which was adjusted as necessary. The IR, area of necrotic spots, and DI were quantified based on the outcomes derived from the ImageJ analyses.

### 2.9 Data statistics

In this study, the t-test, analysis of variance (ANOVA), and least significant difference (LSD) methods were employed to assess the significance of the NA. Additionally, the chi-square test was utilized to evaluate the significance of the IR, while the Shapiro-Wilk test was applied to verify the normal distribution of the numbers of poplar clones across different resistance levels.

## 3 Results

### 3.1 Stem canker pathogen *V. sordida* induces necrotic symptoms on poplar leaves

In this study, both the control (WA inoculation) and *V. sordida* inoculation induced necrotic spots around the pricking sites on poplar leaves at 5 DAI (Figure 2A-B). The necrotic spots in the control were non-expanding, with their average areas measuring 1.5±0.5 mm^2^ (Figure 2A, 2C-F). Conversely, pathogen inoculation resulted in enlarged necrotic spots on poplar leaves (Figure 2B, 2G-K). No hyphal structures were observed under a stereomicroscope on the surfaces of the control poplars. In contrast, hyphal structures were noted on the pathogen-inoculated poplars (Figure 2G-I), and even some pycnidial structures were observed (Figure 2J-K), with some extruding milky white spores noted on the leaves of certain hybrid poplar clones such as B246 (Figure 2K), B73, B14, B18, B76, B181, and B1. In severely diseased clones, such as clone B246, almost all five necrotic spots overlapped. Using the NA on the control poplar leaves as a reference, this study defined NA > 2.0 mm^2^ as the quantitative criterion for disease onset on the pathogen-inoculated poplar leaves (Table 2).

### 4.2 The influence of leaf ontogeny on disease severity

This study examined how leaf ontogeny affects disease severity using the 5^th^ to 7^th^ newly matured (upper) leaves and the 18^th^ to 20^th^ mature (lower) leaves of *P. euramericana* cv. ’Bofeng 3’. As illustrated in Figure 4A, at 5 DAI, the disease severity in the upper leaves was significantly lower than in the lower leaves (chi-square test, n=60, *p*<0.05). The percentage of disease incidence (P) for the upper and lower leaves was 28.33% and 70.00%, respectively. After excluding the non-diseased inoculation points, the results indicated that the area of necrotic spots on the upper leaves was also significantly smaller than that on the lower leaves (2.49±0.13 mm^2^ (n=17) versus 2.94±0.13 mm^2^ (n=42); t-test, *p*<0.05). Furthermore, the DI for the upper and lower leaves were 5.67 and 14.67, respectively (Figure 3A).

**Figure 3:**
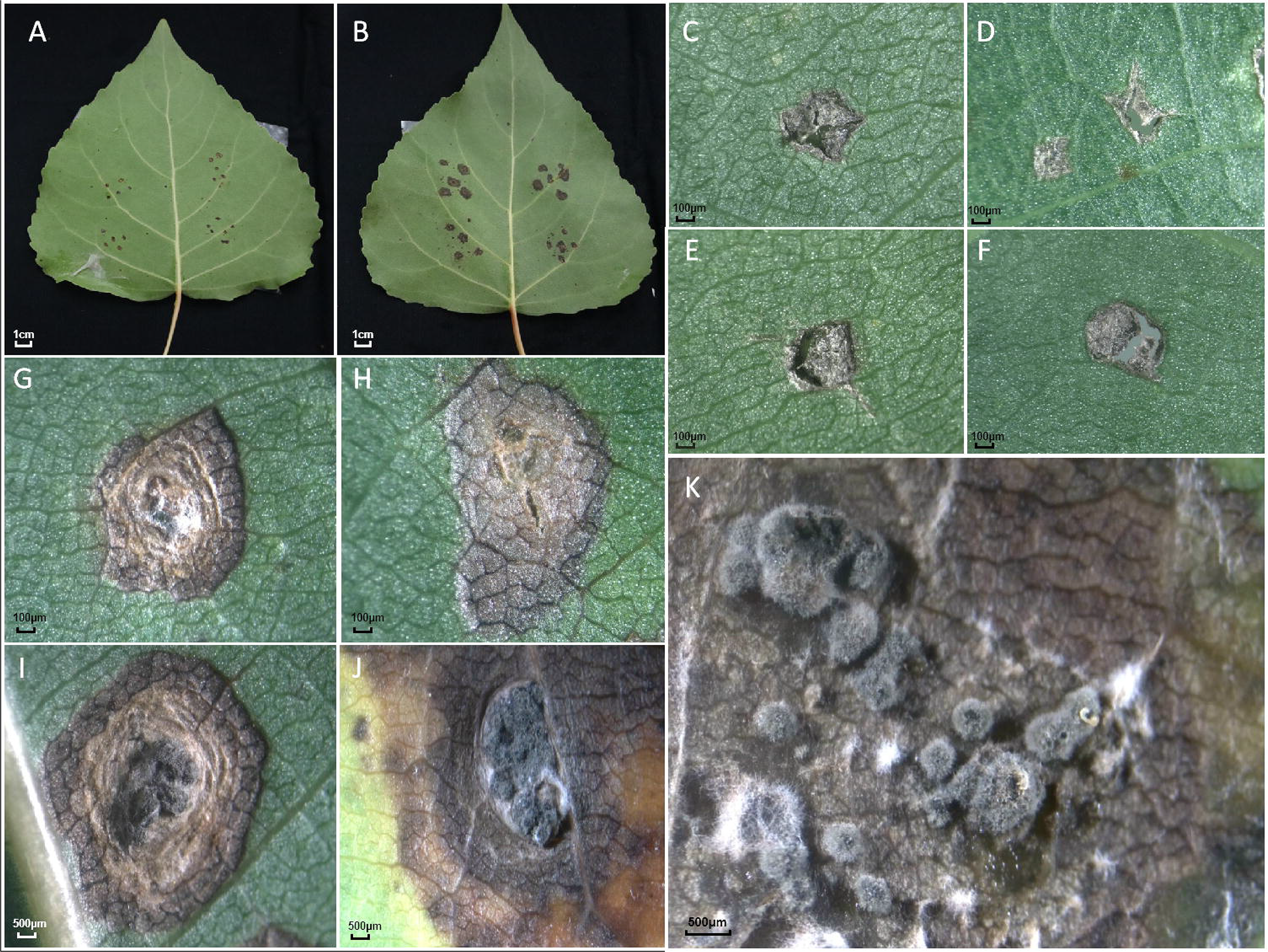
Necrotic symptoms on poplar leaves induced by *V. sordida* isolate CZC using a novel leaf inoculation method in this study. (A) Unextended necrotic spots (1.5±0.5 mm^2^) on poplar leaves inoculated with WA medium. (B) Extended necrotic spots (more than 2.0 mm^2^) on leaves inoculated with *V. sordida* CZC. (C-F) Necrotic spots on leaves inoculated with PDA, at 5 DAI. (G-I) Necrotic spots on *V. sordida* CZC-inoculated leaves, at 5 DAI. (J-K) Formation of fungal hyphae and pycnidial structures on highly susceptible hybrid clone B246, at 5 DAI.

**Figure 4:**
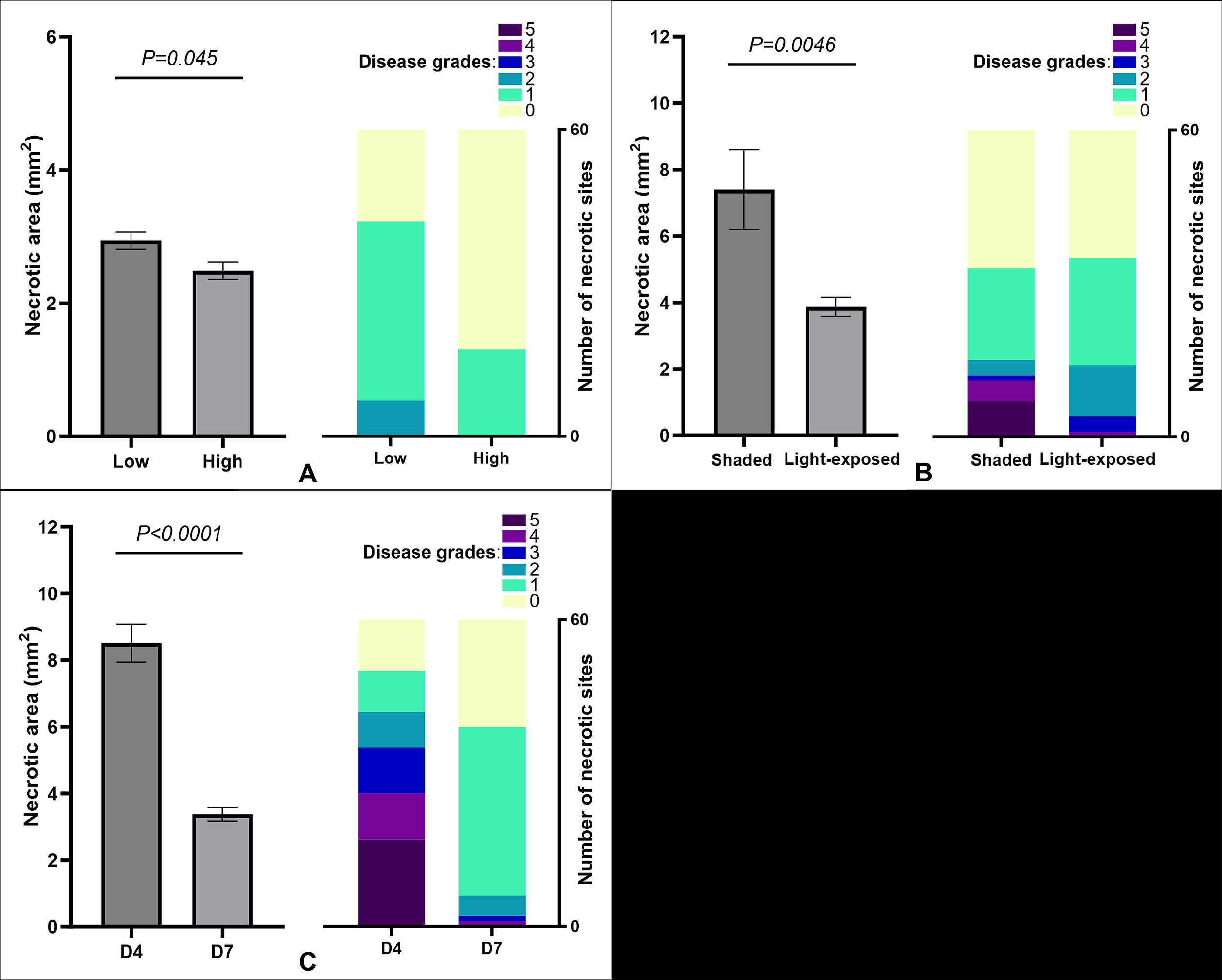
Effects of leaf position (A), light condition (B), and culture age of mycelium inoculants (C) on the necrotic average area and 60 necrotic sites severity of leaf disease induced by *V. sordida* isolate CZC using the leaf inoculation method.

### 3.3 The influence of light condition of leaves on disease severity in leaves

In this study, the 5^th^ to 7^th^ leaves from three light-exposed branches (exposed to sunlight throughout the day) and three shaded branches (not exposed to direct sunlight) of the same 2-year-old saplings were utilized to assess the impact of light conditions on disease severity. The results indicated no significant differences in the IR between the light-exposed and shaded leaves (IRlight- exposed 58.33% vs. IRshaded 55.00%; chi-square test, n=60, *p*>0.05). However, the findings suggested that shading increased disease severity in poplar leaves: the average NA of the light-exposed and shaded leaves was 3.87±0.29 mm^2^ (n=35) and 7.40±1.20 mm^2^ (n=33), respectively (t-test, *p*<0.05). Additionally, the DI for light-exposed leaves (DIlight-exposed) was 18.00, compared to 26.00 for shaded leaves (DIshaded) (Figure 4B).

### 3.4 The influence of culture time of fungal inoculum on disease severity in leaves

Results indicated that the culture time of the mycelium inoculum significantly influenced disease development in poplar leaves. The IR induced by 4-day-cultured *V. sordida* mycelium (IR4day) was 83.33%, which was significantly higher than the rate induced by 7-day-cultured mycelium (IR7day, 65.00%) (chi-square test, n=60, *p*<0.05). This suggests that juvenile pathogen inoculums exhibited higher pathogenicity than their older counterparts. After excluding the non-diseased inoculation points, it was found that the NA induced by 4-day-cultured mycelium plugs (NA4day=8.51±0.57 mm^2^, n=50) was significantly larger than that caused by 7-day-cultured mycelium plugs (NA7day=3.37±0.20 mm^2^, n=39) (t-test, *p*<0.05). Furthermore, DI4day and DI7day, were 50.67 and 13.33, respectively (Figure 4C).

### 3.5 *In vivo* inoculation of stem canker pathogen *V. sordida* on leaves reveals resistance diversity in poplar hybrid clones

In this study, each hybrid clone produced 60 prick sites, and collectively, all 48 clones produced 2,880 sites. After excluding the inefficient inoculation sites (where the mycelium plugs moved away or dried within 1-3 DAI), a total of 2,715 effective inoculation data points were obtained, representing 94.27% of the planned inoculation dataset. By adopting a criterion where the NA > 2.0 mm^2^ indicates the occurrence of disease on poplar leaves, a total of 2,145 inoculation sites exhibited definitive necrotic features, confirming an actual leaf disease incidence of 79.00% across 40 poplar hybrid clones.

Results revealed a remarkable range of resistance diversity to the *Valsa* canker pathogen among the 48 poplar hybrid clones, thanks to the innovative application of a novel leaf inoculation method. As depicted in Table S3, the IR among the 48 poplar clones varied from 31.67% (poplar clone B133) to 100.00% (clones B1 and B104) (Figure 5A). The size of the NA ranged from 2.72±0.13 mm^2^ (clone B168) to 15.28±1.26 mm^2^ (clone B14) (Figure 5B), and the DI varied from 6.67 (clone B168) to 83.67 (clones B246 and B135) (Figure 5C).

**Figure 5:**
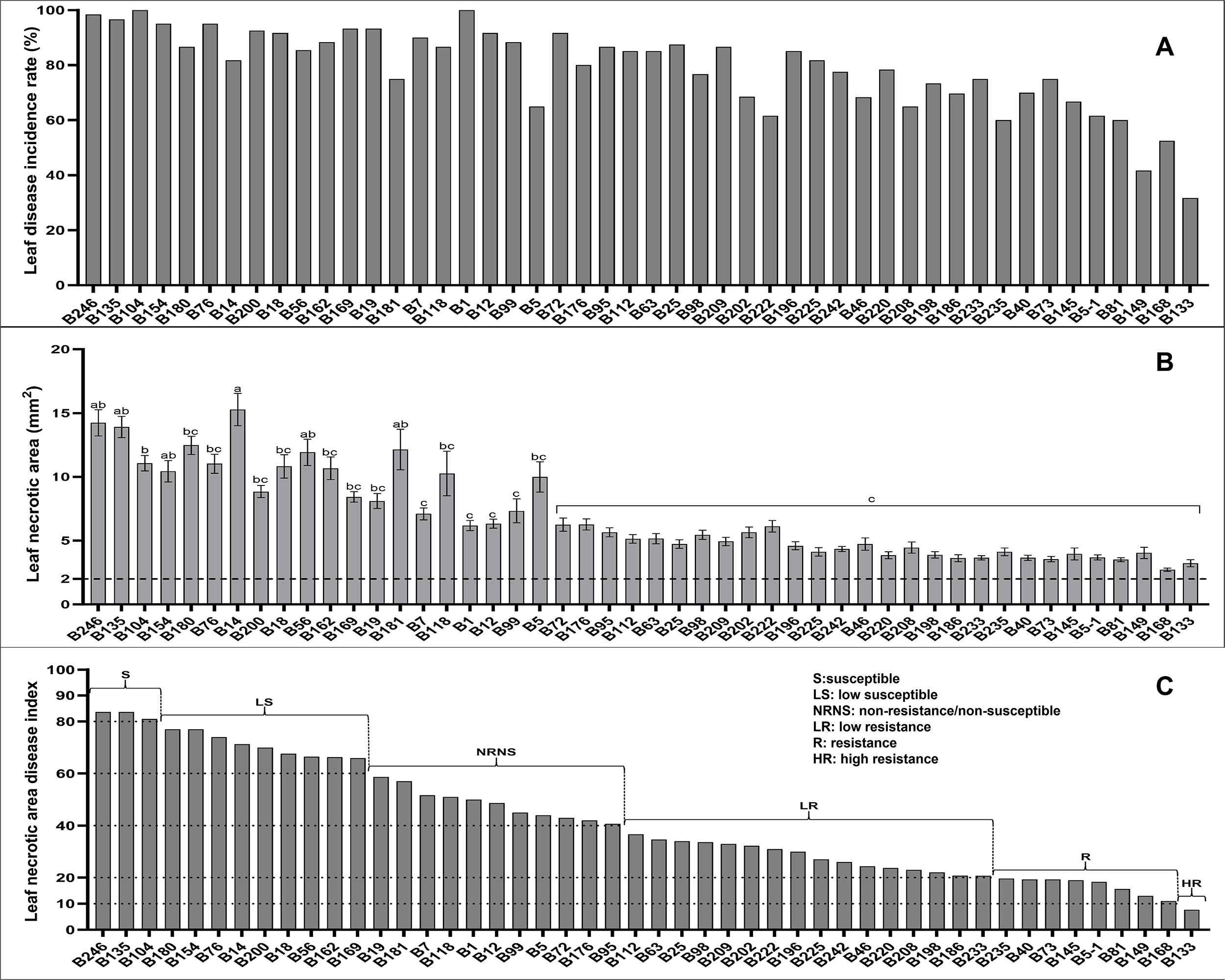
Leaf disease incidence rate (A), leaf average necrotic area (B), and leaf necrotic area disease index (C) from 48 poplar hybrid clones inoculated with *V. sordida* isolate CZC using the leaf inoculation method.

This study employed the Shapiro-Wilk test to evaluate the distribution of resistance levels among 48 poplar clones. Although the p-value from the Shapiro-Wilk test did not support a normal distribution, the absolute kurtosis from the normality test was 1.05 (<10.0), and the absolute skewness was 0.44 (<3.0), suggesting that the resistance distribution among these clones can generally be accepted as normal (Kline, 2023). This conclusion is also supported by the P-P plot analysis (Figure 6).

**Figure 6:**
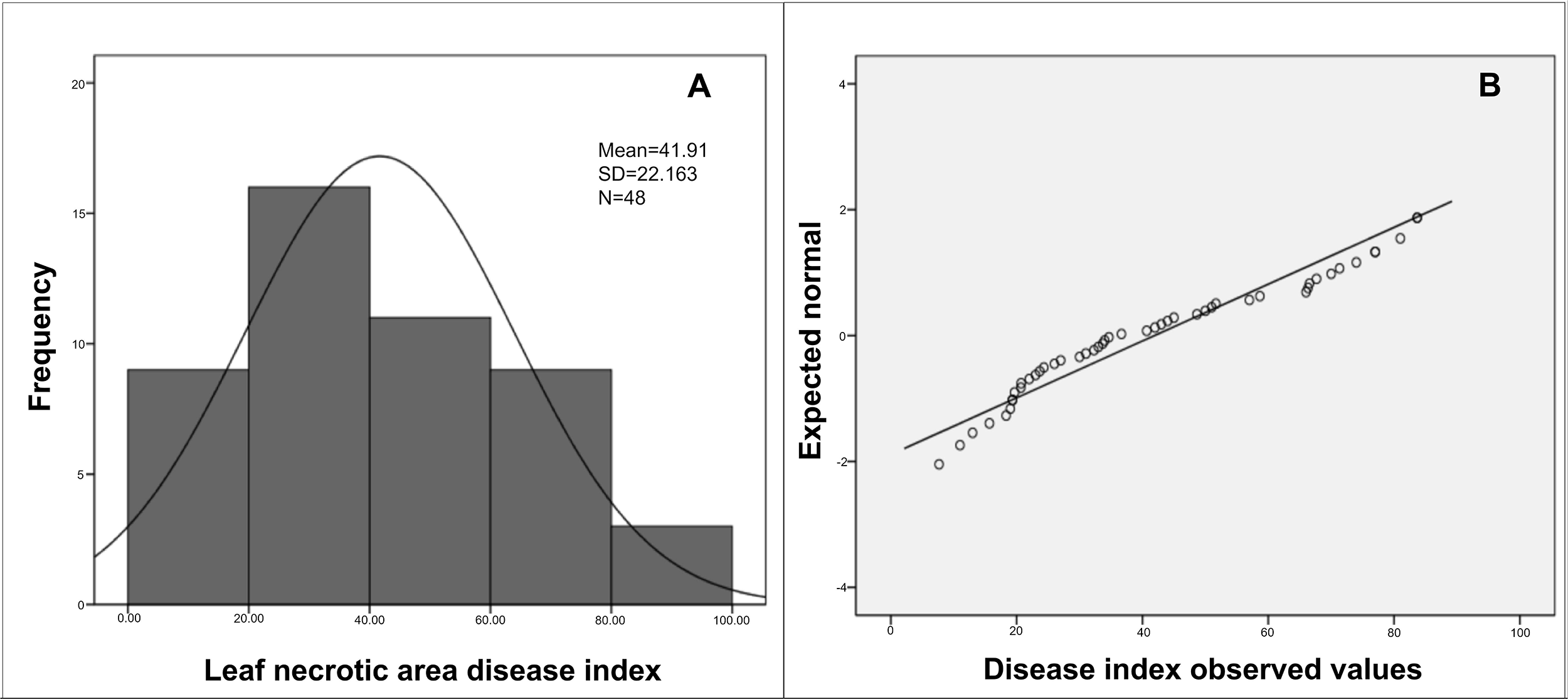
Distribution of resistance among 48 hybrid poplar clones using the leaf inoculation method, showing a generally normal distribution. The standard distribution model (A) and the P-P plot of the clones (B).

According to the DI, this study revealed a broad spectrum of disease resistance among all 48 poplar hybrid clones, ranging from high resistance to susceptibility (Figure 5C). For instance, clone B133 was identified as highly resistant (HR), clones B168 and seven others as resistant (R), while clones B135, B246, and B104 were susceptible (S) to the *Valsa* canker pathogen CZC, as determined by these leaf assessment methods (Table S3).

### 3.6 Verification of the leaf inoculation method through *in vitro* stem segment inoculation

As indicated in Table 3, the IR among 15 poplar hybrid clones ranged from 18.00% (B133) to 98.00% (B246). The Stem necrotic time disease index (DI*stem-time*) varied from 15.71 (B133) to 92.86 (B246), and the stem necrotic area disease index (DI*stem-area*) ranged from 5.00 (B133) to 78.00 (B246) (Figure 7A-C). The values and rankings of leaf necrotic area disease index (DI*leaf-area*), stem disease incidence rate (IR*stem*), DI*stem-time*, and DI*stem-area* were compared to evaluate the consistency between leaf and stem inoculation methods. According to the data presented in Table 3, except for clone B145 in the R/HR group and clone B76 in the susceptible/least susceptible (S/LS) group, the resistance rankings of the remaining eight poplar clones assessed by both leaf and stem inoculation methods were consistent. Notably, the rankings of the most resistant clone, B133, and the least resistant clone, B246, were identical across both methods. Two-dimensional graphs plotted for the DI*leaf-area* versus the DI*stem-time*, the IR*stem,* the DI*stem-area* are displayed in Figure 7D-F, respectively. Simple linear regression analysis yielded R^2^ values of 0.84, 0.79, and 0.52, there is an interrelationship between the results of leaf inoculation method and results of stem segment inoculation method (Zou *et al*., 2003). Additionally, the clustering of different poplar clones based on the leaf inoculation method largely coincided with the results of poplar disease resistance grading obtained through the stem segment inoculation method. Therefore, it can be concluded that there is a significant consistency in the results of disease resistance testing for different poplar clones between the leaf inoculation method and the stem segment inoculation method. These findings support the leaf inoculation method as a viable alternative for identifying pathotypes and assessing resistance in poplar breeding and tree pathological research.

**Figure 7:**
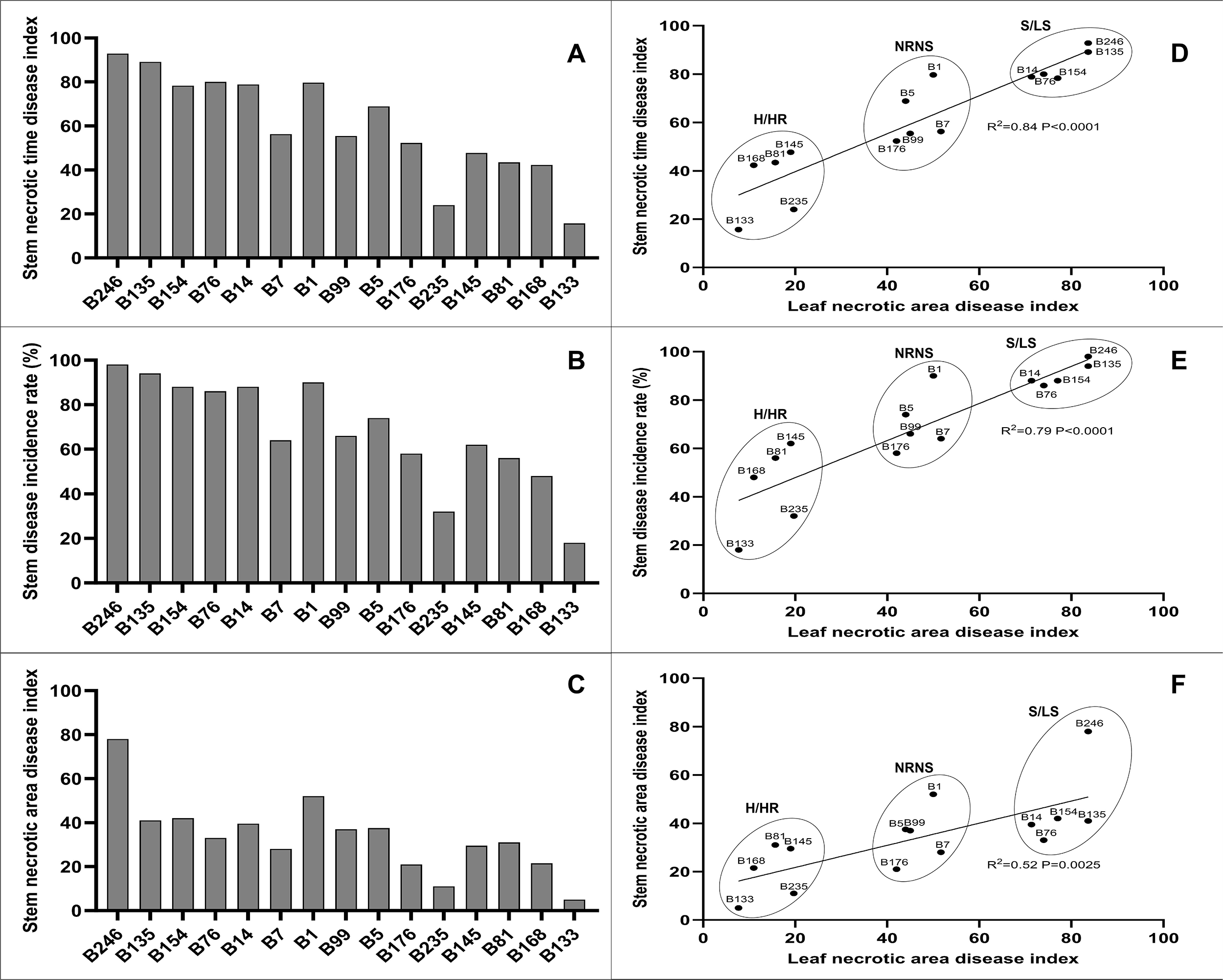
Stem necrotic time disease index (DIstem-time) (A), stem disease incidence rate (IRstem) (B), stem necrotic area disease index (DIstem-area) (C), two-dimensional representation of DIstem-time and leaf necrotic area disease index (DIleaf-area) (D), two-dimensional representation of IRstem and DIleaf-area (E), two-dimensional representation of DIstem-area and DIleaf-area (E) of the stems of 15 poplar hybrid clones were inoculated by poplar stem canker *V. sordida* isolate CZC using the stem inoculation method.

**Table 3.** Disease index of 15 poplar hybrid clones, detected using *in vivo* leaf inoculation and *in vitro* stem segment inoculation methods. Rankings are based on results from the leaf inoculation method

### 3.7 *In vivo* leaf inoculation reveals diversity of pathogenicity in stem canker pathogen *B. dothidea*

The findings of this study hold significant practical implications for the field of plant pathology. The pathogenicity of 12 *B. dothidea* isolates on the poplar hybrid ’Bofeng 3’ varied substantially. The isolate SD47 induced the highest NA in poplar leaves, measuring 6.57±0.41 mm^2^, while the NAs caused by isolates SD60 and CZ1218 were 4.53±0.68 mm^2^ and 3.20±0.19 mm^2^, respectively. Although the differences between SD47 and SD60, as well as between SD60 and CZ1218, were not statistically significant, the NA induced by SD47 was significantly greater than that by CZ1218 (ANOVA, *p*<0.05) (Figure 8). Furthermore, the NAs of nine isolates, including CZ1070c, CZ1070b, CZ1010, and CZ1055, were smaller than 2.0 mm^2^, indicating their low pathogenicity on ’Bofeng 3’ (Table 4). The quantitative criteria used in this study confirmed the low pathogenicity of these nine isolates on the poplar hybrid ’Bofeng 3’ (Table 4). These results underscore the utility of the leaf inoculation method in assessing pathogen pathogenicity, presenting a valuable tool for future research and disease management strategies.

**Figure 8:**
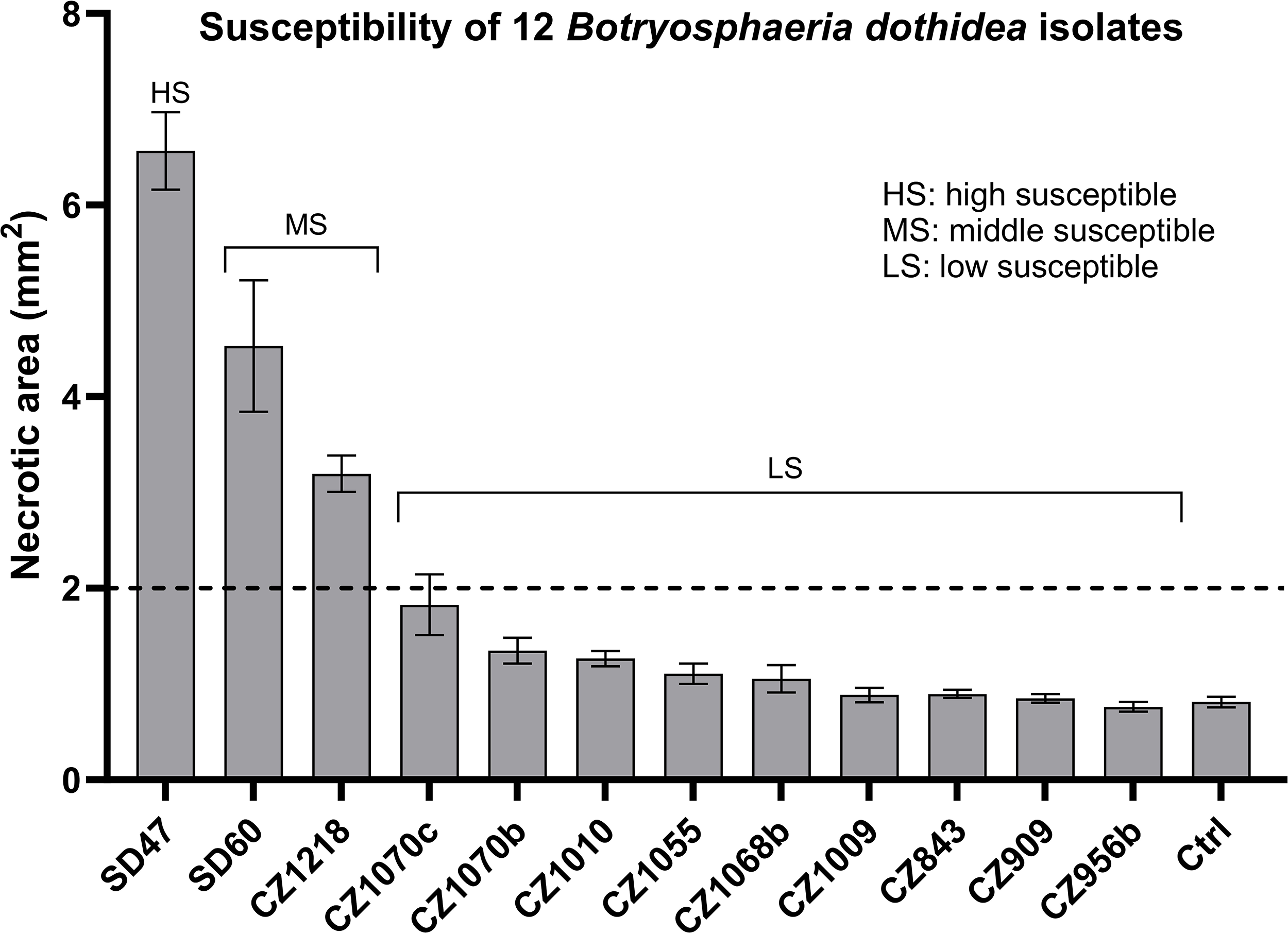
The average necrotic area in the inoculated leaves of "Bofeng 3" poplar against 12 *B. dothidea* isolates.

**Table 4.** Leaf average necrotic areas on ’Bofeng 3’ poplar leaves inoculated with 12 *B. dothidea* isolates.

## 4 Discussion

Selection based on phenotypic traits is a critical step in hybrid breeding. Tree species often exhibit characteristics such as large crowns, slow growth, and lengthy maturity periods, which necessitate more time, space, and human resources for phenotypic acquisition in the context of tree stem canker diseases compared to herbaceous diseases and tree leaf diseases. Additionally, the dispersion of necrotic spots across various heights on the branches further complicates and increases the complexity of acquiring phenotypes for stem canker diseases, making the process more tedious and challenging than for leaf diseases.

Numerous fungal pathogens, including *Valsa*, *Botryosphaeria*, *Leucostoma*, and *Septoria*, contribute to canker diseases affecting poplar branches (Hayova, 1998; Dunnell & LeBoldus, 2017). *Valsa* and *Botryosphaeria* are considered the most significant fungal pathogens in Chinese poplar plantations (Zhang & Luo, 2003). Research has demonstrated that stem canker pathogens can infect host leaves or reside within leaves as endophytes, offering a viable pathway to detect resistance to stem canker disease through leaf inoculation (Marsberg *et al*., 2017; Feau *et al*., 2010; LeBoldus *et al*., 2010; Dunnell & LeBoldus, 2017; Zheng *et al*., 2017; Wei *et al*., 2010). However, the consistency of the leaf inoculation method compared to the stem inoculation method remains under debate. Studies by Wei *et al*. (2010) and findings from the current study on *Valsa* canker disease have shown consistency between these two methods, suggesting that leaf inoculation is an effective technique for selecting resistant clones or for conducting assays of pathogenic differentiation within the pathogen population. Nevertheless, there was no correlation in disease severity between leaf spot and stem canker in the *poplar-Septoria* pathosystem (Dunnell & LeBoldus, 2017).

The severity of disease following leaf inoculation correlates directly with leaf resistance. The association between leaf age (and position on branches) and resistance has been well-documented for over a century (Gusberti *et al*., 2013). This phenomenon, known as ontogenic or age-related resistance, manifests specifically as leaf-stage resistance in leaf tissues (Asalf *et al*., 2014; Dunnell & LeBoldus, 2017; Xu *et al*., 2018). In physiological studies, due to their active and stable physiological performance, the newly matured leaves at the top of branches or saplings (typically the 5^th^ to 7^th^ leaves) are considered ideal for researching photosynthesis, chlorophyll fluorescence, stomatal movement, water status, physiology, biochemistry, and gene expression regulation in tree species (Zhao *et al*., 2017; Xing *et al*., 2019; Li *et al*., 2019; Shen *et al*., 2023; Li *et al*., 2024). In the case of powdery mildew in strawberries and *Malus-Venturia* interactions, leaf resistance rapidly increases from the first unfurled and expanding leaf to the lower leaves, peaking at the 5^th^ to 6^th^ matured expanding leaves (Asalf *et al*., 2014; Gessler & Stumm, 1984; MacHardy *et al*., 1996). In the *Malus-V. ceratosperma* system, the inoculated leaves are identified as "the fully extended leaves at the top of the stems" (Wei *et al*., 2010). This study demonstrated that the resistance of the 5^th^ to 7^th^ leaves (always newly matured) is significantly higher than that of the 18^th^ to 20^th^ leaves (with 32-35 leaves growing on the 1-year-old saplings). Furthermore, the resistance findings from the 5^th^ to 7^th^ leaves were consistent with those from the excised stem segments (Table 3, Figure 7D-F). However, despite differences in disease type (*Valsa* canker vs. *Septoria* canker) and inoculation method (wounding mycelium vs. non-wounding spore inoculation), variations in leaf position (5^th^-6^th^ newly matured leaves vs. the five middle leaves in branches of 3 or 4-month-old hybrid poplar clones) might be the primary cause of inconsistency between leaf and stem resistance in the poplar-*Septoria* pathosystem (Dunnell & LeBoldus, 2017). Therefore, the analysis suggests that the top 5^th^ to 7^th^ leaves are optimal inoculation materials for evaluating and selecting resistant clones in poplar breeding.

Solar UVB radiation serves as a positive modulator of plant defense (Ballaré, 2014). Evidence from various plant-pathogen systems indicates that shading exacerbates infections by various pathogens (Roberts & Paul, 2006). Additionally, high population density, often associated with reduced light exposure, significantly augments disease severity in both natural and agricultural systems (Augspurger & Kelly, 1984; Bell *et al*., 2006; Burdon & Chilvers, 1982; Jurke & Fernando, 2008). In this study, the leaf inoculation method was employed to detect resistance in perennial poplars and to assess the pathogenicity of different pathogen isolates using leaves from various branches of the same poplar tree. The light conditions experienced by these leaves vary substantially, particularly between shaded and light-exposed leaves. This variability led to investigating whether such differences could influence pathogen susceptibility. The results confirm that susceptibility to the pathogen CZC increased significantly under shaded conditions in the hybrid poplar ’Bofeng 3’ (Figure 4B). Shading is commonly utilized to facilitate disease onset in controlled environments. For example, in the hybrid-*Septoria* system, branches were covered with black plastic garbage bags, sealed with two pieces of moist paper towels, for 48 hours to enhance spore germination and invasion (LeBoldus *et al*., 2010; Dunnell & LeBoldus, 2017). Similarly, small necrotic spots were observed on light-exposed leaves in these experiments (Figure 4B). Moreover, another study found that high density or shading induced more severe canker disease symptoms on the trunk of 6-year-old *P. alba* var. *Pyramidalis* (Shen *et al*., 2024). Therefore, in assays evaluating poplar resistance or pathogen pathogenicity using the leaf inoculation method, the specific light conditions of the poplar leaves must be carefully considered.

In the Petri dish, the morphology, colors, and growth rates of colonies and mycelia of the pathogens varied during the culture period. Given that plants can exhibit age-related resistance, such as leaf-stage resistance, pathogens may also demonstrate a similar age-related pathogenicity. For instance, the pathogenicity and infection rate of *Dactylaria higginsii* conidia, harvested from 15 days of culturing on PDA, were higher than those from other culture durations (Kadir *et al*., 2011). Conidiospores of three entomopathogenic fungi (*Metarhizium anisopliae*, *Paecilomyces fumosoroseus*, and *Verticillium lecanii*) harvested from young cultures (2–3 days old) on Petri dishes germinated more rapidly than those from older cultures. Conversely, conidia of *Beauveria bassiana* germinated at consistent rates, regardless of the culture age (Richard *et al*., 1994). This study found that the pathogenicity of young cultures (4 days old on PDA medium) of *V. sordida* isolate CZC to poplars was higher than that of older cultures (7 days old). However, some fungal isolates cannot grow to over 9 cm in diameter on a Petri dish in 7 days; for instance, certain *V. sordida* isolates require specific culture conditions to yield more mycelium for inoculation: culture on PDA medium at 28°C in dark conditions for seven days.

Some researchers posit that the most aggressive or virulent strains of a pathogen should be utilized for selecting and testing plant germplasm and breeding lines (Balendres *et al*., 2019). While this strategy may aid in identifying the most resistant clones within a breeding population, it could potentially compromise the entire breeding program if it results in widespread disease or the demise of all progeny, especially if other agronomically important traits are neglected. As illustrated in Figure 6, the distribution of poplar clones across seven different pathogenicity levels of the pathogen *V. sordida* isolate CZC approaches a normal distribution, indicating that *V. sordida* isolate CZC is an optimal test strain for the current hybrid poplar population.

If another isolate of *V. sordida*, with higher pathogenicity than the *V. sordida* isolate CZC, were used, the distribution of disease susceptibility among these 48 poplar clones might exhibit a right- skewed pattern (with most clones susceptible to the pathogen). Conversely, employing an isolate with weaker pathogenicity might result in a left-skewed distribution. Therefore, it is crucial to preselect the optimal pathogen strains in a new breeding program before conducting final resistance assays. The side length of the square mycelium plugs, 1.0-1.2 cm, is an experiential value derived from our preliminary experiments; for assays involving a strain of high pathogenicity, larger plugs may be more effective. Additionally, while three isolates of *B. dothidea* demonstrated noticeable pathogenicity, the other nine fungal isolates showed no or very low pathogenicity towards the hybrid poplar ’Bofeng 3’. This lack of pathogenicity may be attributed to the high disease resistance inherent in the hybrid. Consequently, the pre-selection of poplar clones with suitable resistance levels is essential when assessing large-scale pathogen populations.

This study demonstrated that the poplar stem canker pathogen *V. sordida* can induce lesion symptoms on leaf tissues using a leaf inoculation method. The effectiveness of this method was validated within a small hybrid poplar population. The process is straightforward and easy to implement; a team of three can inoculate over 200 clones in a single day. For the rapid development of disease symptoms, pathotypes can be acquired within 3-5 DAI; consequently, breeders can identify the pathotypes of a large-scale population (1,000 genotypes or more) within one month. When disease stress is limited to brief periods (specifically, removed from the stems at 3-5 DAI), the pathogen minimally impacts the growth and physiology of the inoculated poplars, thus preserving key phenotypic traits such as height, stem diameter, and mass. Therefore, breeders can quickly implement multi-objective breeding strategies that enhance resistance to canker disease while also promoting rapid growth and high yield. Although extensive post-inoculation data are still required to fully support this research, the feasibility of this method has been validated through observation of disease progression on leaves and subsequent stem inoculation. Moving forward, this study plans to apply the leaf inoculation method to a larger population of clonally propagated poplar trees with verified disease resistance differentiation, further confirming the feasibility of this approach.

In summary, the *in vivo* leaf inoculation method provides a rapid, efficient, cost-effective, and high-throughput alternative to the more time-consuming and costly *in vitro* stem segment inoculation method. This technique is especially suitable for large-scale resistance assays of hybrid poplar clones and can be utilized in the selection of resistant seedlings for breeding stem canker diseases, as well as certain leaf diseases, in poplars and other tree species. When integrated with advanced breeding technologies, such as genomic selection, this method enables a deep exploration of breeding resources to identify new resistance-related genes, gene loci, or gene modules (Du *et al*., 2023). Additionally, the leaf inoculation method can be employed for large-scale pathogenicity testing and differentiation assays of both poplar and other tree species afflicted by stem canker or leaf pathogens.

## 6 Author Contributions

Conceptualization, J.Z. and B.Z.; methodology, J.Z. and B.Z.; software, Z.L. and Y.S.; validation, Z.L., Y.S., W.S., and Y.F.; formal analysis, Z.L., Y.S., W.S., and Y.F.; investigation, Z.L., Y.F., and Y.S.; resources, B.Z. and J.Z.; data curation, Z.L., Y.S., and J.F.; writing—original draft preparation,

Z.L. and J.Z.; writing—review and editing, J.Z., Y.Z., Z.L., H.L., and L.P.; visualization, Z.L. and W.S.; supervision, X.S.; project administration, J.Z. and B.Z.; funding acquisition, J.Z. All authors have read and agreed to the published version of the manuscript.

## Acknowledgments

We thank the Zhao lab members for their constructive suggestions and insightful discussions.

## 8 Data Availability Statement

Data will be made available upon request.

**Table S1.** Severity grading criteria for disease onset time in poplar stems

**Table S2.** Severity grading criteria for disease onset area in poplar stems

**Table S3.** Leaf average necrotic areas of 48 hybrid poplar clones inoculated with *V. sordida* isolate CZC.

