## Supplementary material for "A rapid and efficient *in vivo* inoculation method for introducing tree stem canker pathogens onto leaves: Suitable for large-scale assessment of resistance in poplar breeding progeny": Table 1

Table 1. **Hosts and locations of fungal isolates used in this study**.

| Isolates | Pathogen | Host | Location | Latitude and longitude | Citation |
| --- | --- | --- | --- | --- | --- |
| CZC | *Valsa sordida* | *Populus sp.* | Beijing botanical garden, Haidian, Beijing | 40.00°N,116.21°E | Xing *et al.*, 2020 |
| SD47 | *Botryosphaeria dothidea* | *Populus sp.* | Nanzhaolou, Yuncheng, Shandong | 35.44°N,115.90°E | - |
| SD60 | *B. dothidea* | *Populus sp.* | Huanhai Road, Yantai, Shandong | 37.38°N,119.94°E | Wang 2013 |
| CZ1218 | *B. dothidea* | *Populus sp.* | Guanbei, Huayin, Shaanxi | 34.55°N,110.12°E | Wang 2013 |
| CZ1070c | *B. dothidea* | *Populus sp.* | Shenzhou, Hengshui, Hebei | 37.99°N,115.56°E | Wang 2013 |
| CZ1070b | *B. dothidea* | *Populus sp.* | Shenzhou, Hengshui, Hebei | 37.99°N,115.56°E | Wang 2013 |
| CZ843 | *B. dothidea* | *Populus sp.* | Hebei Agricultural University, Baoding, Hebei | 38.82°N,115.44°E | Wang 2013 |
| CZ909 | *B. dothidea* | *Populus sp.* | Dajidian, Baoding, Hebei | 38.82°N,115.39°E | - |
| CZ956b | *B. dothidea* | *Populus sp.* | Tangxian, Baoding, Hebei | 38.93°N,114.78°E | - |
| CZ1009 | *B. dothidea* | *Populus sp.* | Renxian, Xingtai, Hebei | 37.12°N,114.65°E | - |
| CZ1010 | *B. dothidea* | *Populus sp.* | Renxian, Xingtai, Hebei | 37.12°N,114.65°E | Wang 2013 |
| CZ1055 | *B. dothidea* | *P. tomentosa* | Pingshan, Shijiazhuang, Hebei | 38.43°N,113.90°E | Wang 2013 |
| CZ1068b | *B. dothidea* | *Populus sp.* | Shenzhou, Hengshui, Hebei | 37.99°N,115.56°E | Wang 2013 |
