## Supplementary material for "A rapid and efficient *in vivo* inoculation method for introducing tree stem canker pathogens onto leaves: Suitable for large-scale assessment of resistance in poplar breeding progeny": Table 2

Table 2. Severity grading of necrotic symptoms induced by canker pathogens on leaves

| Severity grade | Grading criteria | Level |
| --- | --- | --- |
| No disease | No symptoms, necrotic area 0-2.0mm² | 0 |
| Extremely mild | Necrotic area 2.0-4.0mm² | 1 |
| Mild disease | Necrotic area 4.0-6.0mm² | 2 |
| Moderate disease | Necrotic area 6.0-8.0mm² | 3 |
| Moderately severe disease | Necrotic area 8.0-10.0mm² | 4 |
| Severe disease | Necrotic area greater than 10.0mm² | 5 |
