## Supplementary material for "A rapid and efficient *in vivo* inoculation method for introducing tree stem canker pathogens onto leaves: Suitable for large-scale assessment of resistance in poplar breeding progeny": Table 3

Table 3. **Disease index of 15 poplar hybrid clones, detected using *in vivo* leaf inoculation and *in vitro* stem segment inoculation methods. Rankings are based on results from the leaf inoculation method**

| Poplar clones | Resistance level of poplar clones based on leaf inoculation method | DI_leaf-area_ (ranking) | IR_stem_ (ranking) | DI_stem-time_ (ranking) | DI_stem-area_  (ranking) | Average of ranking in leaves and stems | Average of ranking in stems |
| --- | --- | --- | --- | --- | --- | --- | --- |
| B246 | S | 83.67 (1) | 98.00 (1) | 92.86 (1) | 78.00 (1) | 1.00 | 1.00 |
| B135 | S | 83.67 (2) | 94.00 (2) | 89.14 (2) | 41.00 (4) | 2.50 | 2.67 |
| B154 | LS | 77.00 (3) | 88.00 (4) | 78.29 (6) | 42.00 (3) | 4.00 | 4.35 |
| B76 | LS | 74.00 (4) | 86.00 (6) | 80.00 (3) | 33.00 (8) | 5.25 | 5.67 |
| B14 | LS | 71.33 (5) | 88.00 (5) | 78.86 (5) | 39.50 (5) | 5.00 | 5.00 |
| B7 | NRNS | 51.67 (6) | 64.00 (9) | 56.29 (8) | 28.00 (11) | 8.50 | 9.33 |
| B1 | NRNS | 50.00 (7) | 90.00 (3) | 79.71 (4) | 52.00 (2) | 4.00 | 3.00 |
| B99 | NRNS | 45.00 (8) | 66.00 (8) | 55.43 (9) | 37.00 (7) | 8.00 | 8.00 |
| B5 | NRNS | 44.00 (9) | 74.00 (7) | 68.86 (7) | 37.50 (6) | 7.50 | 6.67 |
| B176 | NRNS | 42.00 (10) | 58.00 (11) | 52.29 (10) | 21.00 (13) | 11.00 | 11.33 |
| B235 | R | 19.67 (11) | 32.00 (14) | 24.00 (14) | 11.00 (14) | 13.25 | 14.00 |
| B145 | R | 19.00 (12) | 62.00 (10) | 47.71 (11) | 29.5 (10) | 10.75 | 10.33 |
| B81 | R | 15.67 (13) | 56.00 (12) | 43.43 (12) | 31.00 (9) | 11.50 | 11.00 |
| B168 | R | 11.00 (14) | 48.00 (13) | 42.29 (13) | 21.50 (12) | 13.00 | 12.67 |
| B133 | HR | 7.67 (15) | 18.00 (15) | 15.71 (15) | 5.00 (15) | 15.00 | 15.00 |
