## Supplementary material for "A rapid and efficient *in vivo* inoculation method for introducing tree stem canker pathogens onto leaves: Suitable for large-scale assessment of resistance in poplar breeding progeny": Table 4

Table 4. Leaf average necrotic **areas on 'Bofeng 3' poplar leaves inoculated with 12 *B. dothidea* isolate**s.

|  | Necrotic areas (Mean ± SEM, mm^2^) |  | Necrotic areas (Mean ± SEM, mm^2^) |
| --- | --- | --- | --- |
| SD47 | 6.57±0.41 | CZ909 | 0.85±0.05 |
| SD60 | 4.53±0.68 | CZ956b | 0.76±0.05 |
| CZ1218 | 3.20±0.19 | CZ1009 | 0.89±0.08 |
| CZ1070c | 1.83±0.32 | CZ1010 | 1.27±0.08 |
| CZ1070b | 1.35±0.13 | CZ1055 | 1.11±0.11 |
| CZ843 | 0.90±0.04 | CZ1068b | 1.06±0.14 |
