## Supplementary material for "A rapid and efficient *in vivo* inoculation method for introducing tree stem canker pathogens onto leaves: Suitable for large-scale assessment of resistance in poplar breeding progeny": Table S1

Table S1. Severity grading criteria for disease onset time in poplar stems

| Days after inoculation/d | ≤5 | 5＞～≤10 | 15＞～≤20 | 20＞～≤25 | 25＞～≤30 | 30＞～≤35 | ＞35 |
| --- | --- | --- | --- | --- | --- | --- | --- |
| Grade | 7 | 6 | 4 | 3 | 2 | 1 | 0 |
