## Supplementary material for "A rapid and efficient *in vivo* inoculation method for introducing tree stem canker pathogens onto leaves: Suitable for large-scale assessment of resistance in poplar breeding progeny": Table S2

Table S2. Severity grading criteria for disease onset area in poplar stems

| Area (A)/mm² | ≥11.5 | 7.5≥～＜11.5 | 5.4≥～＜7.5 | 3.1≥～＜5.4 | ＜3.1 |
| --- | --- | --- | --- | --- | --- |
| Grade | 4 | 3 | 2 | 1 | 0 |
