## Supplementary material for "A rapid and efficient *in vivo* inoculation method for introducing tree stem canker pathogens onto leaves: Suitable for large-scale assessment of resistance in poplar breeding progeny": Table S3

Table S3. **Leaf** average **necrotic areas of 48 hybrid poplar clones inoculated with *V. sordida* isolate CZC**.

| Hybrid poplar clone | Incidence rate (%) | Necrotic area (Mean ± SEM, mm^2^) | Disease index (DI_leaf-area_) | Resistance level |
| --- | --- | --- | --- | --- |
| B135 | 96.67 | 13.91±0.83 | 83.67 | S |
| B246 | 98.33 | 14.23±1.03 | 83.67 | S |
| B104 | 100.00 | 11.07±0.60 | 81.00 | S |
| B154 | 95.00 | 10.44±0.84 | 77.00 | LS |
| B180 | 86.67 | 12.47±0.71 | 77.00 | LS |
| B76 | 95.00 | 11.02±0.75 | 74.00 | LS |
| B14 | 81.67 | 15.28±1.26 | 71.33 | LS |
| B200 | 92.50 | 8.84±0.47 | 70.00 | LS |
| B18 | 91.67 | 10.82±0.92 | 67.67 | LS |
| B56 | 85.45 | 11.92±1.03 | 66.55 | LS |
| B162 | 88.33 | 10.67±0.89 | 66.33 | LS |
| B169 | 93.33 | 8.43±0.42 | 66.00 | LS |
| B19 | 93.33 | 8.11±0.58 | 58.67 | NRNS |
| B181 | 75.00 | 12.14±1.58 | 57.00 | NRNS |
| B7 | 90.00 | 7.10±0.47 | 51.67 | NRNS |
| B118 | 86.67 | 10.27±1.75 | 51.00 | NRNS |
| B1 | 100.00 | 6.18±0.40 | 50.00 | NRNS |
| B12 | 91.67 | 6.33±0.35 | 48.67 | NRNS |
| B99 | 88.33 | 7.34±0.93 | 45.00 | NRNS |
| B5 | 65.00 | 9.99±1.19 | 44.00 | NRNS |
| B72 | 91.67 | 6.25±0.52 | 43.00 | NRNS |
| B176 | 80.00 | 6.28±0.44 | 42.00 | NRNS |
| B95 | 86.67 | 5.65±0.35 | 40.67 | NRNS |
| B112 | 85.00 | 5.13±0.34 | 36.67 | LR |
| B63 | 85.00 | 5.15±0.41 | 34.67 | LR |
| B25 | 87.50 | 4.73±0.33 | 34.00 | LR |
| B98 | 76.67 | 5.45±0.36 | 33.67 | LR |
| B209 | 86.67 | 4.93±0.33 | 33.00 | LR |
| B202 | 68.57 | 5.65±0.42 | 32.29 | LR |
| B222 | 61.67 | 6.13±0.45 | 31.00 | LR |
| B196 | 85.00 | 4.60±0.32 | 30.00 | LR |
| B225 | 81.67 | 4.12±0.34 | 27.00 | LR |
| B242 | 77.50 | 4.34±0.20 | 26.00 | LR |
| B46 | 68.33 | 4.74±0.48 | 24.33 | LR |
| B220 | 78.33 | 3.86±0.27 | 23.67 | LR |
| B208 | 65.00 | 4.46±0.44 | 23.00 | LR |
| B198 | 73.33 | 3.88±0.26 | 22.00 | LR |
| B186 | 69.64 | 3.63±0.27 | 20.71 | LR |
| B233 | 75.00 | 3.65±0.17 | 20.67 | LR |
| B235 | 60.00 | 4.12±0.30 | 19.67 | R |
| B40 | 70.00 | 3.66±0.20 | 19.33 | R |
| B73 | 75.00 | 3.55±0.21 | 19.33 | R |
| B145 | 66.67 | 3.96±0.47 | 19.00 | R |
| B5-1 | 61.67 | 3.68±0.20 | 18.33 | R |
| B81 | 60.00 | 3.51±0.15 | 15.67 | R |
| B149 | 41.67 | 4.04±0.44 | 13.00 | R |
| B168 | 52.50 | 2.72±0.13 | 11.00 | R |
| B133 | 31.67 | 3.23±0.27 | 7.67 | HR |
